# Coarse-graining the recognition of a glycolipid by the C-type lectin Mincle receptor

**DOI:** 10.1101/2024.05.17.594645

**Authors:** Maxime Noriega, Robin A. Corey, Evert Haanappel, Pascal Demange, Georges Czaplicki, R. Andrew Atkinson, Matthieu Chavent

## Abstract

Macrophage inducible Ca^2+^-dependent lectin (Mincle) receptor recognizes *Mycobacterium tuberculosis* glycolipids to trigger an immune response. This host membrane receptor is thus a key player in the modulation of the immune response to infection by *M. tuberculosis*, and has emerged as a promising target for the development of new vaccines for tuberculosis. The recent development of the Martini 3 force field for coarse-grained (CG) molecular modeling allow the study of interactions of soluble proteins with small ligands but its use for the study of interactions with lipids remains less explored. Here, we present a refined approach detailing a protocol for modeling such interactions at a CG level using the Martini 3 force field. Using this approach, we studied Mincle and identified critical parameters governing ligand recognition, such as loop flexibility and the regulation of hydrophobic groove formation by calcium ions. In addition, we assessed ligand affinity using free energy perturbation calculations. Our results offer mechanistic insight into the interactions between Mincle and glycolipids, providing a basis for rational design of molecules targeting this type of membrane receptors.

## Introduction

Proteins associated with the cell membrane fulfill numerous important roles in cell biology, *e.g.* in signaling, adhesion, and the detection of pathogens.^1–5^ Membrane proteins are therefore central to infection and disease. Drugging such proteins^6^ to modulate their activity circumvents the need to cross the cell membrane, which often limits a molecule’s efficiency. Indeed, while membrane proteins represent approximately 23% of the human proteome, they are the target of more than 60% of drugs currently on the market.^7,8^ Advances in biomolecular structure elucidation have paved the way for the rationalization of structure-based drug design.^9^ Moreover, by eliminating the need for chemical synthesis, computational methods have proven to be extremely efficient for the design of ligands, allowing screening of large ligand libraries.^10^

Docking-based methods have emerged as key tools for the virtual screening of libraries of drug-like molecules.^10,11^ However, such methods have several limitations: (1) they do not allow adaptation of the target protein structure to a given ligand; (2) the binding site must often be defined in advance by the user; (3) it is difficult to take into account a complex environment such as the cell membrane.^12^ To tackle these limitations, strategies based on molecular dynamics (MD) simulations have emerged.^13,14^ These allow docking results to be confirmed, enable the protein to explore conformational space, and allow the ligand to find a more favorable pose in the binding site.^15,16^ Nevertheless, capturing binding events using all-atom (AA) simulations may be computationally expensive for *in silico* high-throughput drug screening.

Coarse-grained (CG) modeling consists of mapping small sets of atoms onto so-called “beads”, parametrized so as to mimic the chemical properties of the sets, resulting in a considerable reduction in computational cost.^17^ The Martini force field, introduced two decades ago, is widely used for biomolecular CG simulations^18–20^ that allow the sampling of binding events through unbiased simulations.^21,22^ Here, we used such method to study the C-type lectin receptor Mincle, a eukaryotic membrane protein receptor involved in the direct recognition of pathogen-derived glycolipids.^23^ In the context of infection by *Mycobacterium tuberculosis* (*mtb*), Mincle binds trehalose dimycolate (TDM) through the carbohydrate recognition domain (CRD) of its extracellular domain, inducing a strong immune response.^24^ Mincle also binds glycolipids from *Malassezia*,^25^ and *β*-glucosylceramide.^26,27^ Binding of ligands to Mincle activates signal transduction through the phosphorylation of an immunoreceptor tyrosine-based activation motif (ITAM) on the intracellular domain of FcR*γ*, the signaling partner of Mincle.^28^ This phosphorylation activates spleen tyrosine kinase (Syk) and subsequently the CARD9-Bcl10-Malt1 pathway, leading to a pro-inflammatory response^29^ (Fig. 1A).

**Figure 1:**
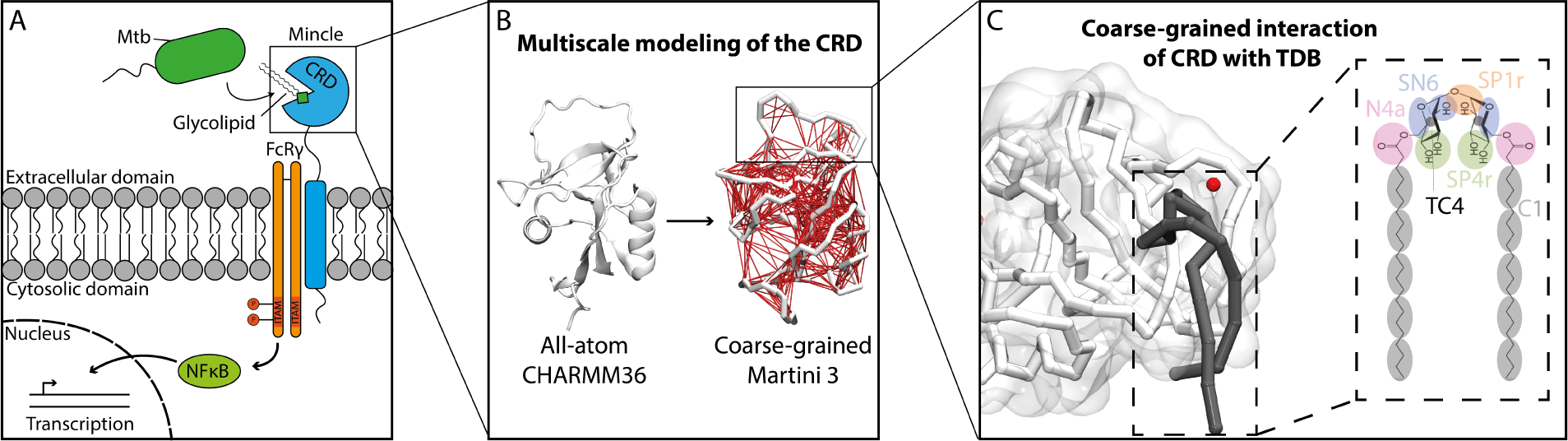
Modeling of Mincle receptor. (A) Schematic representation of Mincle receptor and its partner FcRγ. The binding of glycolipids from *Mycobacterium tuberculosis* (*Mtb*) by the carbohydrate recognition domain (CRD) leads to phosphorylation of the immunoreceptor tyrosine-based activation motif (ITAM) of FcRγ, activating a signaling pathway involving NF-*κ*B that results in the activation of transcription. (B) Coarse-grained modeling of human Mincle CRD. The elastic network is shown in red. (C) Recognition of coarse-grained trehalose dibehenate (TDB) and Ca^2+^ ion (red sphere) by the CRD during simulations.

Targeting Mincle is a promising route to developing adjuvants to strengthen the potency of anti-tuberculosis vaccines, such as BCG.^30^ However, the toxicity of TDM prohibits its use as adjuvant. Trehalose dibehenate (TDB) is a non-toxic synthetic analog of TDM with shorter lipidic chains. It is known to promote a strong immune response that leads to Th1 and Th17 activation.^31–33^

Much structural information has been obtained through X-ray crystallographic studies of the CRDs of bovine and human Mincle.^34–38^ A number of these structures have been solved in the presence of a ligand, including trehalose for bovine Mincle CRD.^34,35^ These studies described the interaction at a canonical binding site comprised of hydrophilic amino acids that form hydrogen bonds with hydroxyl groups of trehalose. The trehalose molecule is oriented such that an attached acyl chain might fit into the adjacent hydrophobic groove formed by Leu172, Val173, Val175, Phe197 and Phe198 of bovine Mincle CRD. This model was further validated by mutagenesis studies.^39^ These investigations also described three binding sites for Ca^2+^ ions, involving: (1) Asp142, Glu146, Asn171, and Asp177; (2) Glu168, Asn170, Glu176, Asn192, and Asp193 and (3) Val116, Asn118, Glu122, and Glu205.

Despite the biological significance of human Mincle, most studies using MD simulations have focused on bovine Mincle CRD.^30,40^ Here, we report CG modeling, using the recently introduced Martini 3 force field,^41^ of the recognition of TDB by human Mincle CRD. Starting from a published crystal structure,^36^ we first parametrized the CG model of human Mincle CRD by fitting to AA simulations (Fig. 1B). We then performed unbiased simulations to observe ligand binding events (Fig. 1C). We employed free energy perturbation (FEP) calculations^42,43^ to further characterize the positioning of the acyl chain of TDB inside the hydrophobic groove as a case study of soluble protein-glycolipid interactions.

## Methods

### System preparation

#### All-atom model of human Mincle CRD

For atomistic simulations, the crystal structure of human Mincle CRD (PDB:3WH3, residues 74-210),^36^ stripped of water molecules and calcium ions, was used as the initial conformation. We deprotonated acidic amino acids since the structure was determined at pH 4 while human Mincle shows higher affinity for its ligands at pH values above 7.^39^ The protein was solvated in a (6.3 nm)^3^ simulation box containing TIP3P water^44^ and 0.10 M NaCl. Additionally, 0, 2 or 3 Ca^2+^ ions were added at random positions, corresponding to a CaCl_2_ concentration of 0 mM, 13 mM and 20 mM, respectively. The system was built using CHARMM-GUI.^45–47^

#### Coarse-grained model of human Mincle CRD

A representative conformation of the CRD, in which the hydrophobic groove, defined by the shortest distances between residue pairs Leu-172/Leu-199 and Val-173/Phe-198, is formed, was extracted from the AA simulation. This structure was converted into a CG model using the martinize2 tool^48^ executed on the Martini database (MAD) server.^49^ The CG model of the protein was solvated in a (6.3 nm)^3^ simulation box with Martini water beads. CaCl_2_ and NaCl were added with the ions at random positions, as for the AA systems. To maintain a well-defined fold, an elastic network was used and refined by calculating per-residue Δ*_RMSF_* compared to AA simulations.

For unbiased simulations in the presence of the ligand, the box size was increased to (10 nm)^3^ with 5 mM CaCl_2_ (corresponding to 3 Ca^2+^ ions) and 0.10 M NaCl. Moreover, we modeled a slightly longer section of human Mincle CRD (residues 59-210) in order to mask a hydrophobic patch. The structure of residues 59-79 was taken from the Alphafold2^50,51^ prediction and we used the concatenation tool of Chimera^52^ to carry out the extension. Simulations with the ligand were performed with the refined model or with conformations in which the long loop was constrained in an open or closed configuration, with a cutoff (RC) of 0.9 nm and a force constant of *K* = 500 kJ.mol*^-^*^1^.nm*^-^*^2^.

#### Ligand preparation

The chemical structure of trehalose dibehenate (TDB) was obtained from the PubChem library (CID:11170611). For AA modeling, force field parameters were generated using the Antechamber tool.^53^ Then, the glycolipid was solvated in a TIP3P water box with 0.15 M NaCl and 5 mM CaCl_2_. For CG modeling, we used the available parameters for trehalose^54^ and typical lipid acyl chain values from Martini 3 force field.^41^ Angles between these two moieties were mapped based on AA simulations.

### Simulation setup

Unless specified otherwise, all MD simulations were performed using the GROMACS 2021.3 package.^55^

#### All-atom simulation of human Mincle CRD

AA simulations of human Mincle CRD were performed using CHARMM36.^56^ The energy of the system was minimized using a steepest descent algorithm^57^ followed by 125 ps equilibration stage at constant volume (NVT) by keeping the backbone restrained. We followed by a set of 3 replicates of a 1 µs production run for each CaCl_2_ concentration, using a leap-frog integrator,^58^ and a 2 fs time-step (see Table 1). We applied the Nosé-Hoover thermostat with a reference temperature of 300 K.^59^ The pressure was maintained isotropically using the Parrinello-Rahman barostat with a reference pressure set to 1.0 bar and a compressibility of 4.5*·*10*^-^*^5^ bar*^-^*^1^. Bond lengths were controlled using the LINCS algorithm.^60^

#### Coarse-grained simulations of human Mincle CRD

CG simulations were carried out using the Martini 3 force field.^41^ The energy of the system was minimized using the steepest descent algorithm. This was followed by a 1000 ps equilibration stage under NVT condition and either a 10 µs production run (for refinement of the elastic network) or 10 sets of 20 µs production runs for simulations with TDB (see Table 1), all using the Parrinello-Rahman barostat^61^ with a reference pressure set to 1.0 bar and a compressibility of 3 *·* 10*^-^*^4^ *bar^-^*^1^. The temperature was held at 300 K using the V-rescale thermostat.^62^

#### TDB simulations

For AA simulation of TDB, the GAFF2 force field^63^ was used. The system was minimized for 20 ps under NVE condition followed by 100 ps for equilibration under NPT condition. AA simulations of the TDB was ran using Amber 2020.^64^ The TDB was then simulated for 200 ns with a reference pressure set to 1.0 bar and temperature was held at 300 K. For CG simulation, the system was first minimized under NVT conditions and equilibrated for 2 ns under NPT conditions. The CG TDB in water was then simulated for 1 µs in equal condition to the AA for 1 µs.

### Simulation analysis

Protein structures were rendered using VMD software and average positions of ligands were computed using the Volmap tool.^65^ Root Mean Square Deviation (RMSD), Root Mean Square Fluctuation (RMSF) and Solvent Accessible Surface Area (SASA) values were calculated using GROMACS commands.^55^ Simulations were analyzed using Tcl/VMD and python scripts. Protein-Ca^2+^/Na^+^ or protein-TDB contacts and were calculated as described previously^42,66^ using cutoff value of 5 Å. Occupancy of Ca^2+^ and Na^+^ ions were calculated using a cutoff of 3 Å because of the proximity between site 1 and 2. Hydrophobic groove formation was defined by computing inter-residue distances (Ile173 - Leu199 and Ala174 - Phe198). Loop amplitude was calculated by measuring the average distance between centers of mass of short and long loops during the course of the simulation, taking the first frame as reference. Binding events between the CRD and TDB were counted if both trehalose moiety and acyl chains were within 5 Å of their respective sites formed by Asn171, Arg183, Asp194 and Ile173, Ala174, Phe198, Leu199 respectively.

#### Free energy perturbation calculation

A bound conformation obtained from unbiased simulations was used as a starting point for FEP calculations. TDB was alchemically transformed either by decoupling one of the two chains or symmetrically shortening the two chains, bead by bead. The chains were linearly decoupled during the calculation along a chemical coordinate, *λ*. Dummy atoms were used to turn off the decoupled beads of the acyl chain. Since the perturbation did not involve charged beads, only Lennard-Jones interactions were turned off. These perturbations used a soft core parameter (*σ* = 0.3).

For each *λ* window, the system was minimized using a steepest descent algorithm, followed by 10 ns of simulation under NPT conditions, using a leap-frog integrator at 300 K, as reported.^43^ The alchemical-analysis package was employed for FEP calculations using the MBAR method on 5 replicates for statistics^67,68^ (see Table 2). ΔΔ*G* values were calculated as described in Figure 4. The GROMACS command `gmx mindist` was used to assess whether the ligand remains bound to the CRD.

## Results and discussion

### Ions recognition by human Mincle CRD

We first ran AA simulations of the CRD domain in a water box containing 0, 2 or 3 Ca^2+^ ions. For simulations containing 2 or 3 Ca^2+^ ions, we observed three main binding sites (Fig. 2A and S1A) as seen in the crystal structures of bovine and human Mincle CRD.^34–36^ These three binding sites are formed by polar and negatively charged amino acids (Fig. 2B) in good agreement with previously determined structures. We observed an additional binding site (ABS) (residues Ser90 and Tyr92) not seen in previous works (Fig. S1A). These binding sites were differently explored by Ca^2+^ ions during the course of the simulations (Fig. 2C and S1B). In particular, site 1 remained less occupied than the two other binding sites during the simulations (Fig. 2A and Fig. S1A, C). This may result from lower specificity for Ca^2+^ ions at this binding site, allowing the binding of other ions. Thus, we also monitored Na^+^ ion binding at these different sites. We observed Na^+^ ions binding to Mincle in a manner similar to Ca^2+^ ions (Fig. 2A and S1A, D). In most cases, no Na^+^ ion was found to interact stably when a Ca^2+^ ion interacted with one of the sites, and reciprocally (Fig. 2C and S1B). These results corroborate the observation of a Na^+^ ion in binding site 1 in a previous study.^34^ As Ca^2+^ and Na^+^ ions interact with the same site in our simulations, we may speculate that there is competition between these two ions at the different sites, with increased specificity for the Ca^2+^ ions.

**Figure 2:**
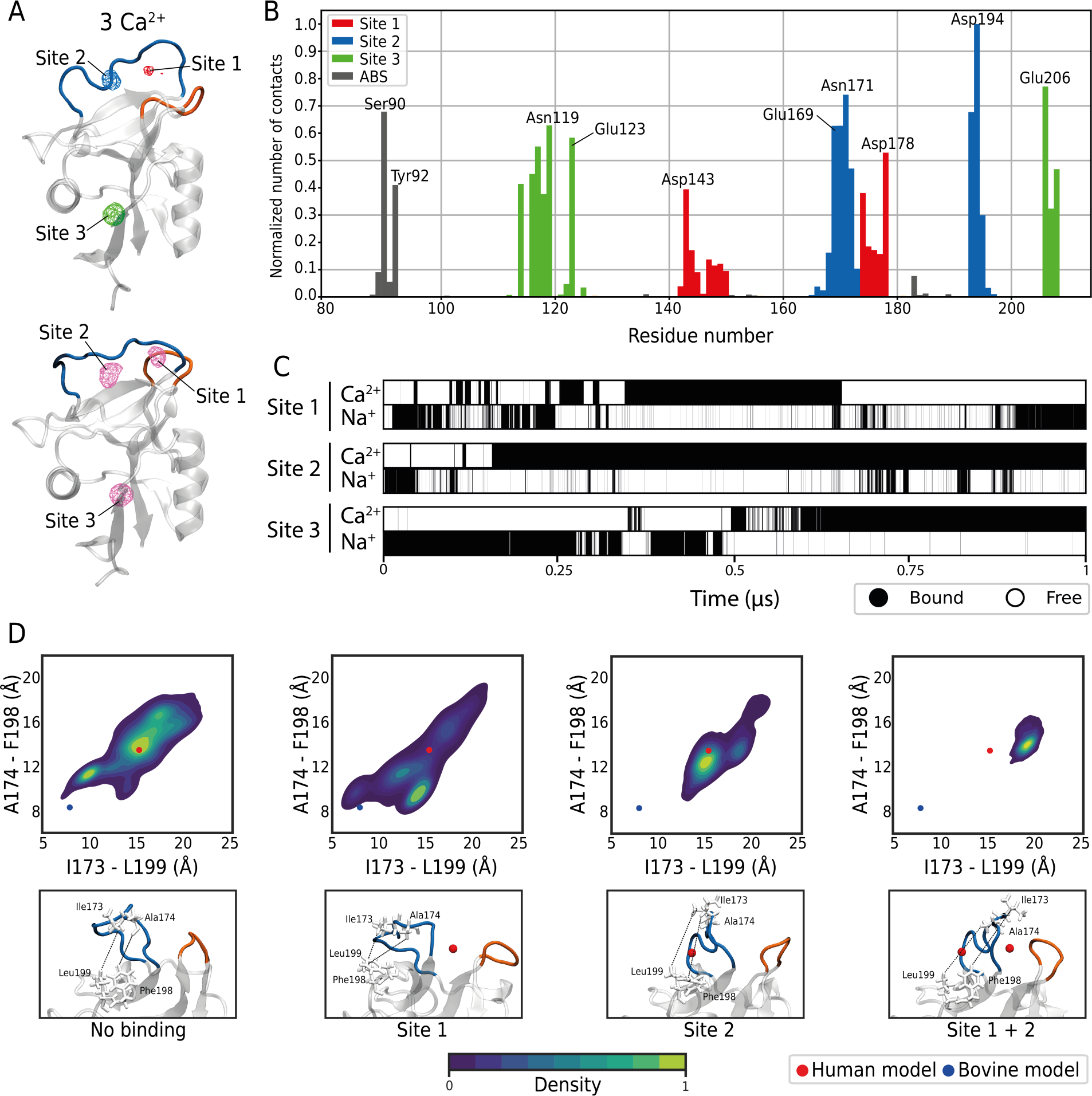
Ion recognition by Mincle receptor. (A) Three-dimensional representation of human Mincle CRD highlighting the long loop (blue) and the short loop (orange) involved in the binding of Ca^2+^ and Na^+^ ions. Average position of Ca^2+^ ions occupying site 1, 2 and 3 are shown in red, blue and green respectively for the simulation containing 3 ions (upper panel, isovalue = 0.1). Average positions of Na^+^ are shown in pink (bottom panel, isovalue = 0.05). These densities were obtained using the volmap tool of VMD. (B) Normalized number of contacts between each amino acids of Mincle and calcium ions. (C) Occupancy of Ca^2+^ and Na^+^ ions at sites 1, 2, 3 during the 1 µs of simulation for one replicate. (D) Average distance between Ile173 - Leu199 and Ala174 - Phe198 according to the occupancy of calcium ions (upper panel). The distances for the bovine and the human crystal structure are represented by blue and red dots respectively. A representative structure is shown for each condition (bottom panel). Ca^2+^ ions are represented by red spheres and side-chains of amino acids forming the hydrophobic groove are represented in white.

### Calcium ions affect the conformation of the loop involved in the ligand binding

Ca^2+^ binding sites 1 and 2 are located near the long loop (Fig. 2A). Residues Ile173 and Ala174 that form the hydrophobic groove, crucial for ligand binding,^30,39^ are located on this loop. Furthermore, this loop was observed to adopt different conformations in bovine (4ZRW) and human (3WH3) Mincle CRD structures.^35,36^ For the human structure (red dot Fig. 2D), hydrophobic residues Ile173 and Ala174 within the long loop are 15 Å apart from the residues Phe198 and Leu199, resulting in an absence of the hydrophobic groove (“loop open”). These distances are about 8 Å for the bovine model (blue dot Fig. 2D), forming the hydrophobic groove (“loop closed”). We therefore investigated conformational changes of the long loop during AA simulations as a function of Ca^2+^ ion binding, starting from the human structure in which the groove is not formed. We extracted, from the 9 replicates of AA simulations, frames for which no calcium ions were bound in sites 1 and 2 (“No binding” Fig. 2D) or for which the sites 1, 2 or both where occupied. For each population, we computed inter-residue distances and compared the loop configuration with that found in bovine and human structures (see methods). In the absence of Ca^2+^ ions in site 2 (Fig. 2D, panels “no binding” and “Site 1”), Ile-173 and Ala-174 approach residues Phe-198 and Leu-199, as seen in the bovine structure, thereby forming the hydrophobic groove. In contrast, binding of Ca^2+^ to site 2 constrains the long loop in an open conformation, thus exploring configurations close to that seen in the human structure and far from that in the bovine structure. Taken together, this may suggest that binding of Ca^2+^ ions to site 2 may restrain the loop far from residues Phe198 and Leu199, avoiding the formation of the hydrophobic groove. On the contrary, these simulations also suggest that the binding of Ca^2+^ ions to site 1 may enable formation of the hydrophobic groove in order to accomodate correctly the acyl chain of glycolipids. These results may explain previous observations where the long loop of bovine Mincle CRD was observed to be open^34^ in the absence of Ca^2+^ in site 1 and closed^35^ in the presence of Ca^2+^ in this site. Similarly, in the C-type lectin receptor DC-SIGN, the long loop was seen to be in a closed conformation when Ca^2+^ ions were bound near to site 1.^69^ In our simulations, both short and long loops appeared flexible (Fig. 2D) allowing the opening of a cleft between the long and the short loop (Fig. S2A and B). These observations may help designing new compounds targeting non-canonical binding sites. Indeed, recent works have targeted a site near the long loop for the C-type lectin receptor Langerin.^70,71^

### Coarse-grained parameterization of human Mincle CRD

To investigate biophysical processes on the time-scale of microseconds, CG MD simulations are commonly used.^72^ CG modeling of proteins requires the use of structural constraints, based on an elastic network, to maintain the secondary structures of the protein.^73–75^ Since the flexibility of the long and short loops are important for both glycolipid recognition and ion binding, we optimized the elastic network based on AA simulations. We first extracted a representative conformation of the CRD from our AA simulations in which the hydrophobic groove is formed. We then converted this conformation into a CG model (Fig. 3A). The underlying elastic network is defined by a set of elastic bonds connecting backbone beads that are separated by a distance lower than a predefined cutoff value (RC).^73^ We tested different RC values to reproduce the intrinsic flexibility of the CRD (Fig. S3). The use of a large RC value (1.0 nm) resulted in an overall CRD flexibility matching that observed in AA simulations, except for the long and short loops which were clearly more flexible in AA simulations (Fig. S3A). Decreasing the RC value increased the flexibility of the loops but introduced too much backbone variability for the remainder of the CRD (Fig. S3A and B). We combined these findings to optimize iteratively the CG model, resulting in an elastic network defined by a RC value of 1.0 nm coupled to a force constant of *K* = 600 *kJ.mol^-^*^1^*.nm^-^*^2^ except for the long and the short loops where RC values of 0.9 and 0.8 respectively coupled to a force constant of *K* = 300 *kJ.mol^-^*^1^*.nm^-^*^2^ were applied. This resulted in flexibility of the CG model in good agreement with AA simulations (Fig. 3B). In particular, the amplitude of motions of the two loops was in better agreement with the AA simulations than before refinement (Fig. 3C and D).

**Figure 3:**
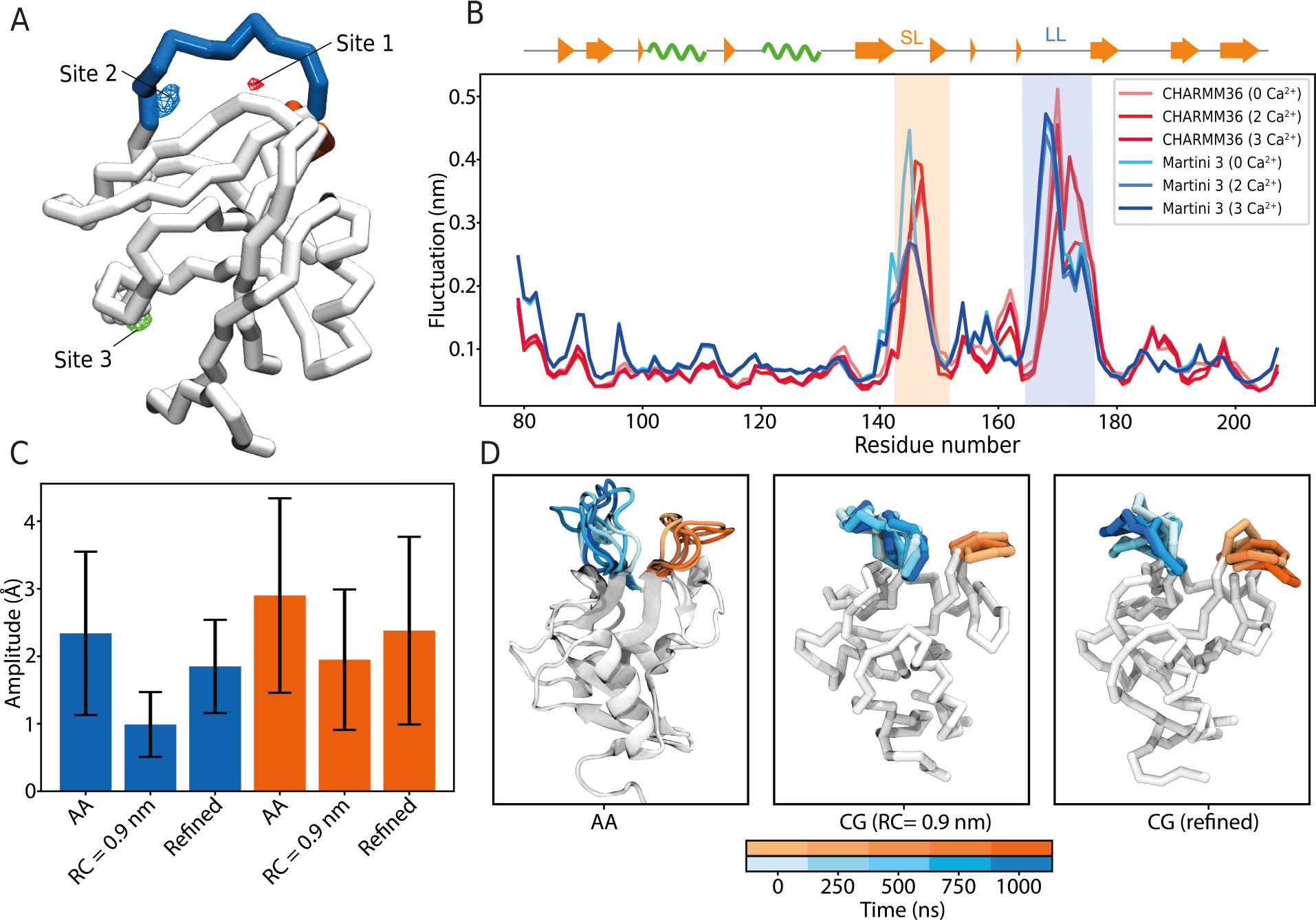
Refinement of the elastic network of the CG model of human Mincle CRD. (A) Binding of Ca^2+^ ions to the refined coarse-grained model of Mincle CRD for the simulation containing 3 Ca^2+^ ions. Long and short loops are shown in blue and orange respectively. Average position of Ca^2+^ ions bound to sites 1, 2 or 3 are shown in red, blue and green respectively. (B) RMSF comparison between AA (red) and CG (blue) simulations. *β*-strands are represented by orange arrows and *↵*-helices by green spirals. (C) Differences in amplitude of motions of the long and short loops (blue and orange, respectively) for AA and CG simulations using an RC value of 0.9 nm or with the optimized model. (D) Example of loop motion for one replicate highlighting the higher flexibility of the refined model compared to the model with an global fixed RC of 0.9 nm.

In our case, it was mandatory that the CG model recover these important binding sites for the protein function. Despite the difference in size between an AA model of an ion and its equivalent CG bead, we were able to identify the main binding site of calcium ions (*i.e.* site 2) with the non-refined models (Fig. S4A). The refined model recovered the site 1 in presence of 3 calcium ions (Fig. 3A). The site 3 was also observed for an RC value of 0.9 nm and 1.0 nm with 3 Ca^2+^ ions or with our refined model in both calcium conditions (Fig. 3A, Fig. S4A, B and C). More generally, it is interesting to notice the ability of Martini 3 CG model to allow the identification of ion binding sites, often crucial in many biological processes.^76^

### Loop flexibility increases glycolipid binding

TDB is composed of a trehalose moiety and two C22 acyl chains linked in 6 and 6’ (Fig. 1C and Fig. S5A). In order to systematize the use of validated parameters, we used topology deposited in the Martini database^49^ corresponding to the trehalose moiety defined by Grünewald *et al.*^77^ and canonical parameters for the lipid acyl chains.^41^ Bond lengths between trehalose and acyl chains were defined based on the monosialodihexosylganglioside (GM3) distances.^77^ Compared to a single trehalose, this link reduces the polarity of the molecule. To take this into account, we replaced the SP1r beads (B2 and B6 position, see Fig. S5A) by SN6 beads. To refine angles parameters, we performed AA simulations of TDB in water and fitted CG values (Fig. S5B). We also compared Solvent Accessible Surface Area (SASA) between AA and CG models. TDB adopts multiple conformations in AA simulations in water resulting in a polydisperse value of the SASA, while the SASA values for the CG model range between 12 and 14 nm^2^ (Fig. S5C and D). This difference is due to stronger interactions between acyl chains in our CG model.

We randomly positioned the TDB CG model in a 10x10x10 nm^3^ box centered on human Mincle CRD containing 5 mM CaCl_2_ and 100 mM NaCl (Fig. 4A). First, we performed simulations with the refined model with the loop fully flexible (see Fig. 3). Then, as controls, we either used the crystal structure (3WH3) in which the long loop is in an open conformation (*i.e.* absence of hydrophobic groove) or a conformation extracted from AA simulations in which the long loop is in closed conformation (*i.e.* presence of hydrophobic groove). For each condition, we performed a set of 10x20 µs CG simulations. While binding events were observed both closed conformation and refined model (Fig. 4B and movie 1), TDB sampled a region near the expected binding site for the trehalose and the hydrophobic groove only for the refined model (Fig. S6A and B). No binding event was observed for CG simulations starting from the crystal structure of human Mincle CRD (*i.e.* open conformation, see Fig. 4B). These results indicate that the loop needs to be flexible to form the hydrophobic groove and accommodate the ligand.

**Figure 4:**
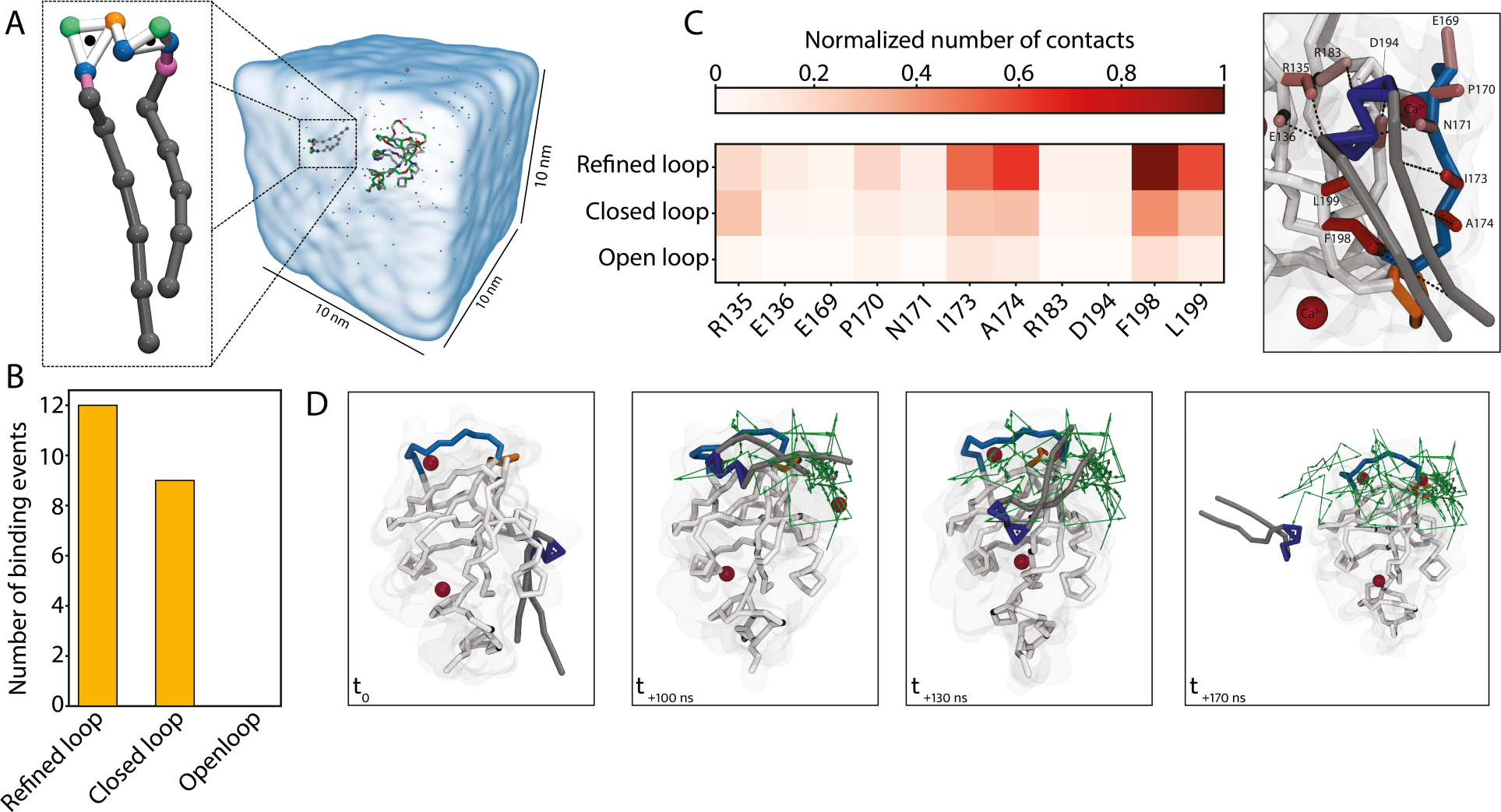
Recognition of TDB by human Mincle CRD. (A) Simulation box showing the CG TDB. (B) Number of binding events between the canonical binding site of thd CRD and the TDB for the refined elastic network or with the long loop maintained in an “open” (corresponding to the crystal structure) or “closed” conformation. (C) Normalized number of contacts of amino acids involved in TDB recognition (left panel). Representative structure of the binding of TDB to human Mincle CRD. Side-chains of amino acids involved in binding are colored according to the color scale. (D) Representation of a binding event between human Mincle CRD and TDB. The center of geometry of the trehalose moiety is tracked by green arrows. Long and short loops are represented in blue and orange, respectively, Ca^2+^ ions are shown in red, the trehalose moiety in dark blue and lipidic chains in gray.

For interactions with the canonical binding site, Glu169, Asn171 and Asp194 were involved in the coordination of Ca^2+^ ions and the binding of the trehalose moiety of TDB (Fig. 4C), as seen in previous studies,^34–37,39^ while amino acids Ile173, Ala174, Phe198 and Leu199, forming the hydrophobic groove, interacted with the acyl chains (Fig. 4C, D).

During these simulations, two other binding sites were identified involving mainly hydrophobic residues. The first site is composed of Pro81, Leu82, Phe93, Phe94, Ala110, Met111, Gly112 and Val208. The second is composed of Phe126, Tyr129 and Pro132 (Fig. S6A, B and C). Binding outside the canonical binding site raises questions about their possible roles. In the context of full-length Mincle receptor, the CRD is attached to the transmembrane domain and may lie near the membrane surface (Fig. 1A). These new binding sites may serve as additional interaction points with the host membrane lipids. Recently, protein/protein interactions involving bovine Mincle CRD were proposed to be mediated by acylated ligand.^38^ These additional binding sites may also help driving such protein interactions.

### Lipid length modulates binding affinity

To better characterize the binding of TDB to human Mincle CRD, we performed free energy perturbation (FEP) calculations based on a canonical binding pose of TDB found during the simulation. We either alchemically transformed the TDB into trehalose monobehenate (TMB) or reduced the length of both lipid chains bead by bead. We also transformed the TDB into two alkyl chains by replacing the trehalose moiety with dummy atoms. We estimated the free energy of binding of these modified TDB structures relative to the whole TDB. To do so, we modeled the full free energy cycle to pass from TDB to shortened structures (Fig. 5A). We performed these calculations in the presence or absence of a Ca^2+^ ion bound to site 2 (Fig. 5B). Removing one acyl chain or reducing its length progressively decreased the binding affinity with a ΔΔ*G* varying between +0.5 and +5.5 kJ.mol*^-^*^1^. Removing one or the other acyl chain gave two different ΔΔ*G* values (*i.e.* +2.5 to +3.5 kJ.mol*^-^*^1^). This energy difference can be explained by the initial position of each acyl chain as chain 1 was positioned closer to the hydrophobic groove. Decoupling the trehalose resulted in a ΔΔ*G* of 4.8 and 3.9 kJ.mol*^-^*^1^ in presence or absence of Ca^2+^ ions respectively. ΔΔ*G* values were not affected by removing the Ca^2+^ ion when the acyl chains were removed. However, according to these FEP calculation, Ca^2+^ ions may stabilize the binding of the trehalose (Fig. 5B). Finally, complete removal of both acyl chains led to unbinding of the trehalose (Fig. S7).

**Figure 5:**
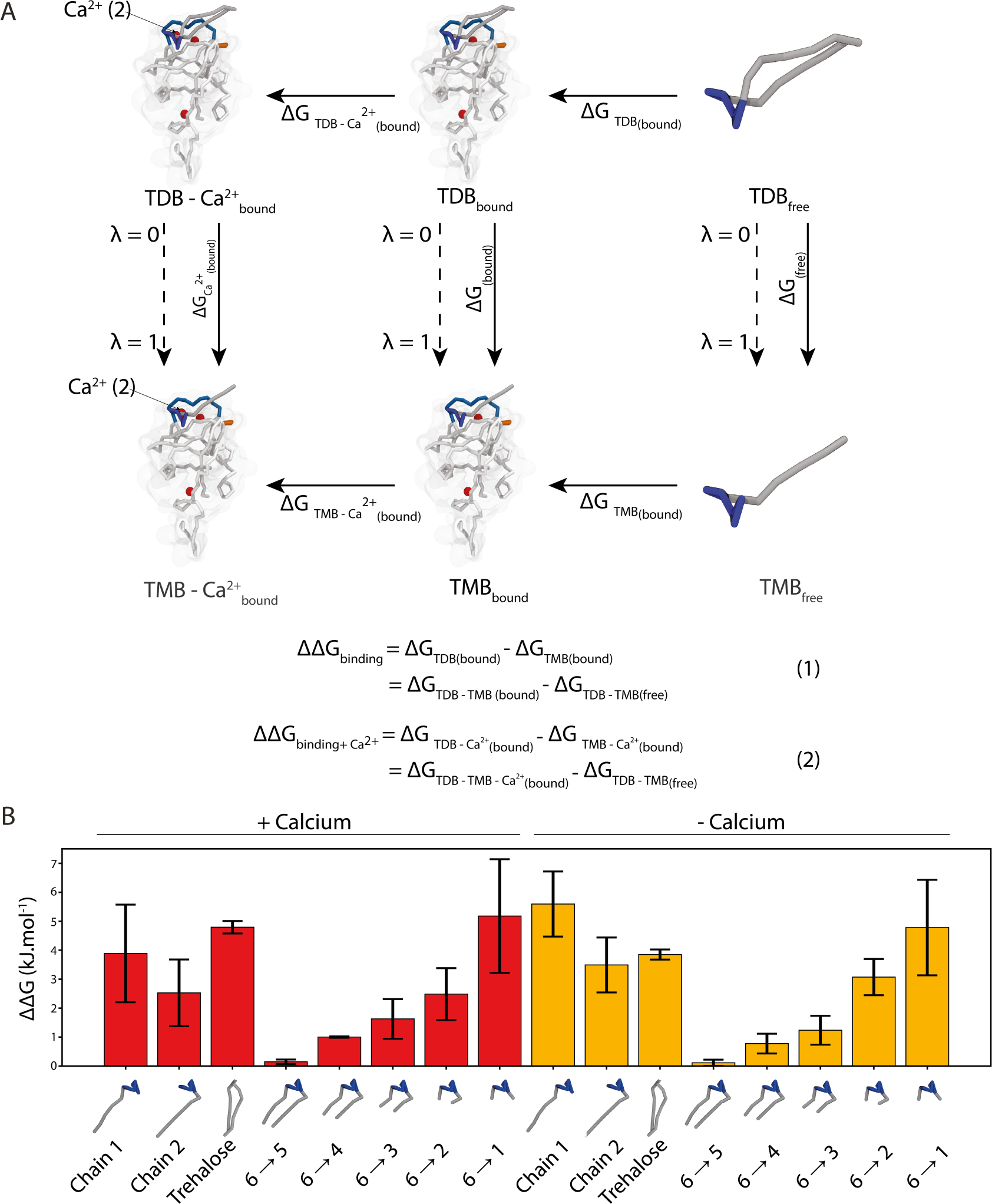
Free energy perturbation. (A) Thermodynamic cycle used for FEP calculations. Bound TDB is shown in blue (trehalose) and gray (acyl chains) and the protein in white. Calcium ions are represented by red spheres. The dashed lines represent the chemical coordinate for the decoupling (*λ*). The bound TDB (2 acyl chains) is alchemically transformed into TMB (1 acyl chain) either in presence or absence of the Ca^2+^ ion bound to site 2 of the CRD. Similar calculations are performed without the protein to calculate ΔΔ*G* values as described by the equations (1) and (2). (B) Means and standard errors of free energy calculation from five replicates. The first or second chain are fully decoupled or the length of both chains is gradually reduced. The trehalose moiety is also decoupled for these calculations. The results are shown in presence or absence of the bound Ca^2+^ ion.

As seen during the unbiased simulation, the interaction with the glycolipid is mainly driven by hydrophobic interactions. FEP calculations confirmed the importance of the length of the acyl chain. Feinberg *et al.* suggested that at least six carbons are needed for the ligand to be correctly accommodated in the hydrophobic groove.^34^ In our case, decoupling 1, 2 or 3 beads did not lead to a considerable loss of free energy of interaction. Decoupling of 4 beads (corresponding to 8 carbon atoms) or more led to a loss of free energy, that was more significant, in good agreement with previous observations.^34^ The decoupling of a full chain (corresponding to the TMB) also led to an important loss of affinity as seen by Decout *et al.*^30^ Together, these data are in line with previous experimental studies.^30,34,39^ Although the interaction is mainly driven by hydrophobic interactions, the loss of binding energy of the ligand from the protein when the trehalose is decoupled suggests that both moieties are important for glycolipid recognition. It is noteworthy that interactions of such magnitude (around 5 kJ.mol*^-^*^1^) were observed for interactions of lipids with membrane proteins using the Martini force field.^43,78^

## Conclusion

Targeting membrane proteins is of interest because it avoids the need to develop compounds that are able to cross the cell membrane. The Martini force field is particularly well suited to study such proteins however, until recently, it was not used routinely to model protein-ligand interactions.^21^ Here, we have assessed a challenging test case for such a force field by studying interactions between the soluble Carbohydrate Recognition Domain of human Mincle with a glycolipid. To do so, we parameterized a CG model of human Mincle using atomistic simulations and carefully modeled loops flexibility, important for glycolipid recognition. We also parameterized a CG model of the glycolpipid TDB and observed multiple binding events to the CRD at various positions, including the canonical binding site of Mincle. We finally assessed which parts of the ligand were the most important for binding using FEP calculations and identified the acyl chains as major contributors to this interaction. Thus, our approach constitutes a good proof of concept of the utility of CG force field for the study of molecules binding to lectins. As the Martini force field is now better suited to perform high-throughput screening of small molecules, the use of CG modeling of lectin-like receptors may serve as an initial phase for high-throughput screening.

## Data availability

Scripts used to analyze MD simulations, AA and CG models of the CRD and the TDB and trajectories are available on Zenodo: https://zenodo.org/records/11186155.

