## Supplementary material for "Coarse-graining the recognition of a glycolipid by the C-type lectin Mincle receptor": SI

Supporting Table 1: **Summary of production simulation.** For the study of the CRD, AA and CG systems were composed of 23K and 2K particles respectively. For simulation in presence of the ligand, the CG systems were composed of 9K particles

| System | Number of $Ca^{2+}$ | Replicates × simulation time |
|---|---|---|
| AA CRD | 0 | 3 × 1 µs |
| | 2 | 3 × 1 µs |
| | 3 | 3 × 1 µs |
| CG CRD | 0 | 4 × 10 µs |
| | 2 | 4 × 10 µs |
| | 3 | 4 × 10 µs |
| CG CRD + TDB (refined) | 3 | 10 × 20 µs |
| CG CRD + TDB (closed) | 3 | 10 × 20 µs |
| CG CRD + TDB (open) | 3 | 10 × 20 µs |

Supporting Table 2: **Summary of simulation for FEP calculation.** As for production, the CG systems were composed of 9K particles. Each calculation were replicated 5 times for statistics, each of the 21 windows were simulated for 10 ns.

| System | Mutation | Simulation times | Replicates × Nb. of windows |
|---|---|---|---|
| + Ca2+ | Chain 1 | 10 ns | 5 × 21 |
| | Chain 2 | | |
| | Trehalose | | |
| | 6 to 5 | | |
| | 6 to 4 | | |
| | 6 to 3 | | |
| | 6 to 2 | | |
| | 6 to 1 | | |
| - Ca2+ | Chain 1 | 10 ns | 5 × 21 |
| | Chain 2 | | |
| | Trehalose | | |
| | 6 to 5 | | |
| | 6 to 4 | | |
| | 6 to 3 | | |
| | 6 to 2 | | |
| | 6 to 1 | | |

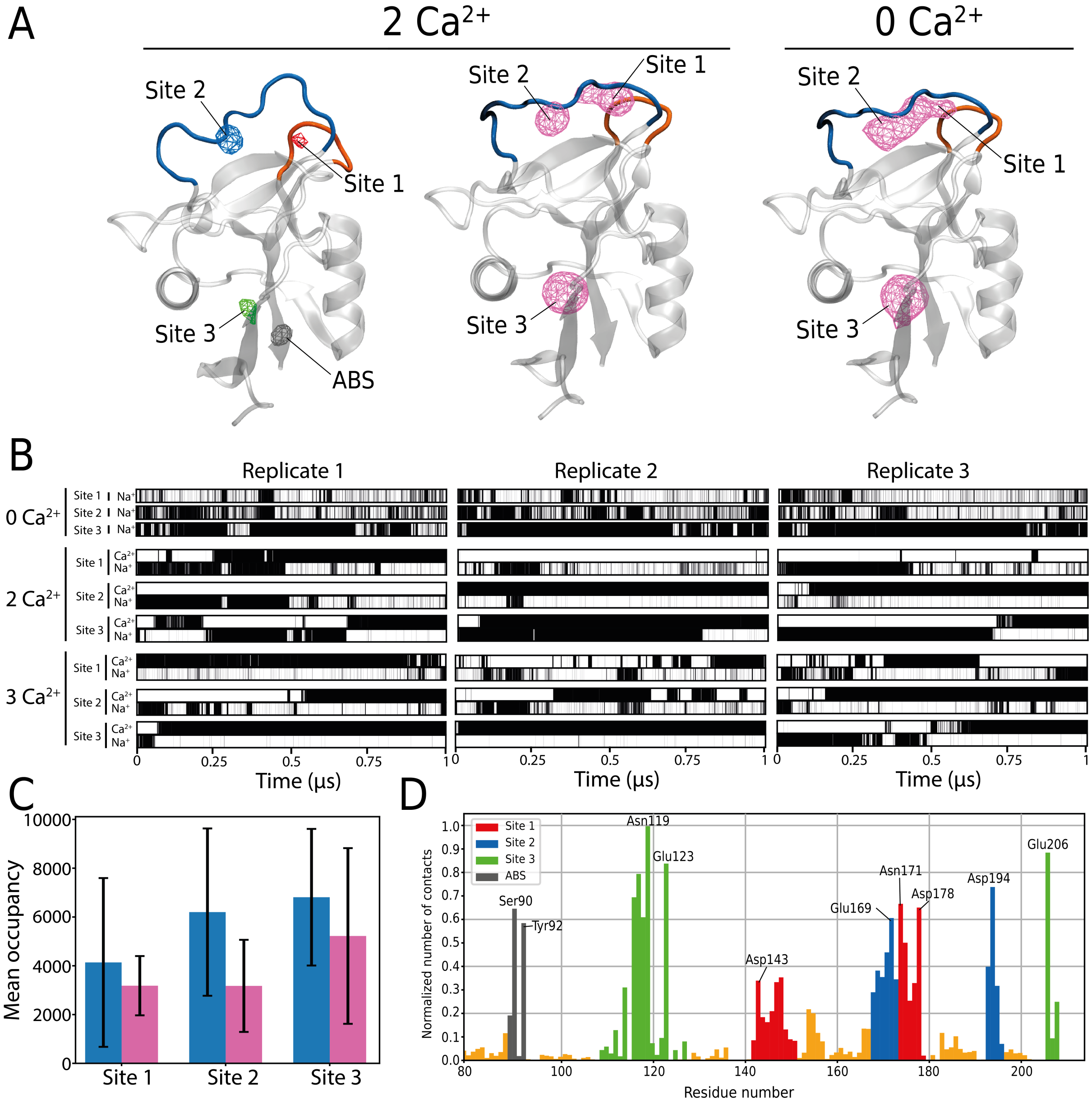


Supporting Figure 1: **Binding of $Ca^{2+}$ and $Na^{+}$ ions by human Mincle CRD** (A) Three-dimensional representation of human Mincle CRD highlighting the long loop (blue) and the short loop (orange) involved in the chelation of $Ca^{2+}$ ions. Average position of $Ca^{2+}$ ions occupying site 1, 2 and 3 are shown in red, blue, and green, respectively for simulations containing 2 calcium ions. An Additional Binding Site (ABS) is shown in gray (isovalue = 0.1). The average position of $Na^{+}$ ions with 0 or 2 calcium ions in the box is shown in pink (isovalue = 0.05). These densities were obtained using the volmap tool of VMD. (B) Occupancy of $Ca^{2+}$ and $Na^{+}$ ions at sites 1, 2, 3 during the 1 µs of simulation for all replicates. Orange bars represent residues that are not associated with any specific site. (C) Mean occupancy of $Ca^{2+}$ (blue) or $Na^{+}$ (pink) ions for sites 1, 2 and 3. (D) Normalized number of contacts between residues of Mincle and $Na^{+}$ ions.

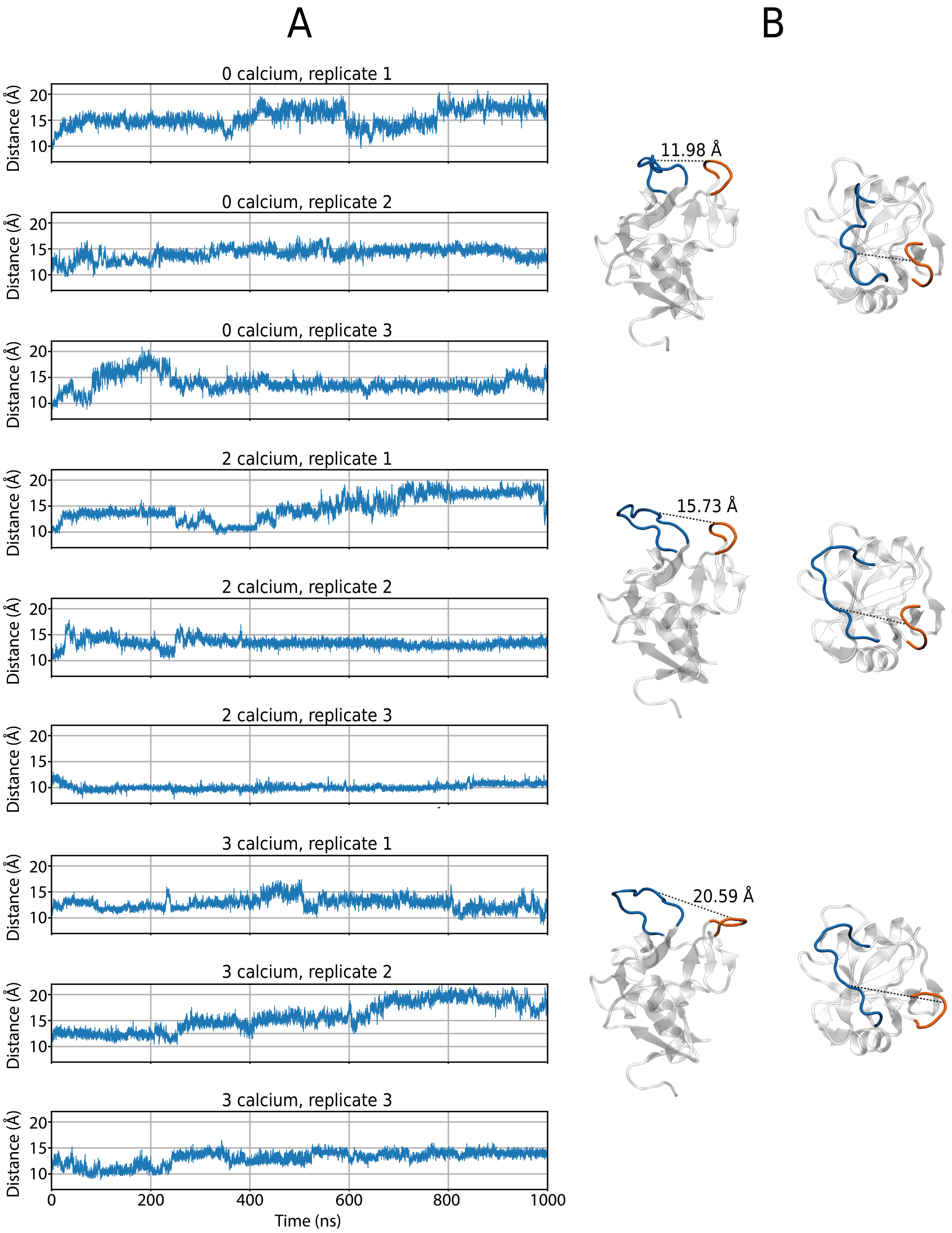


Supporting Figure 2: **Long and short loops flexibility of human Mincle CRD** (A) Distances between the center of mass of both short and long loops in every conditions. (B) Side and top view of human Mincle CRD with its two loops at short (top panel), intermediate (middle panel) and long distances (bottom panel).

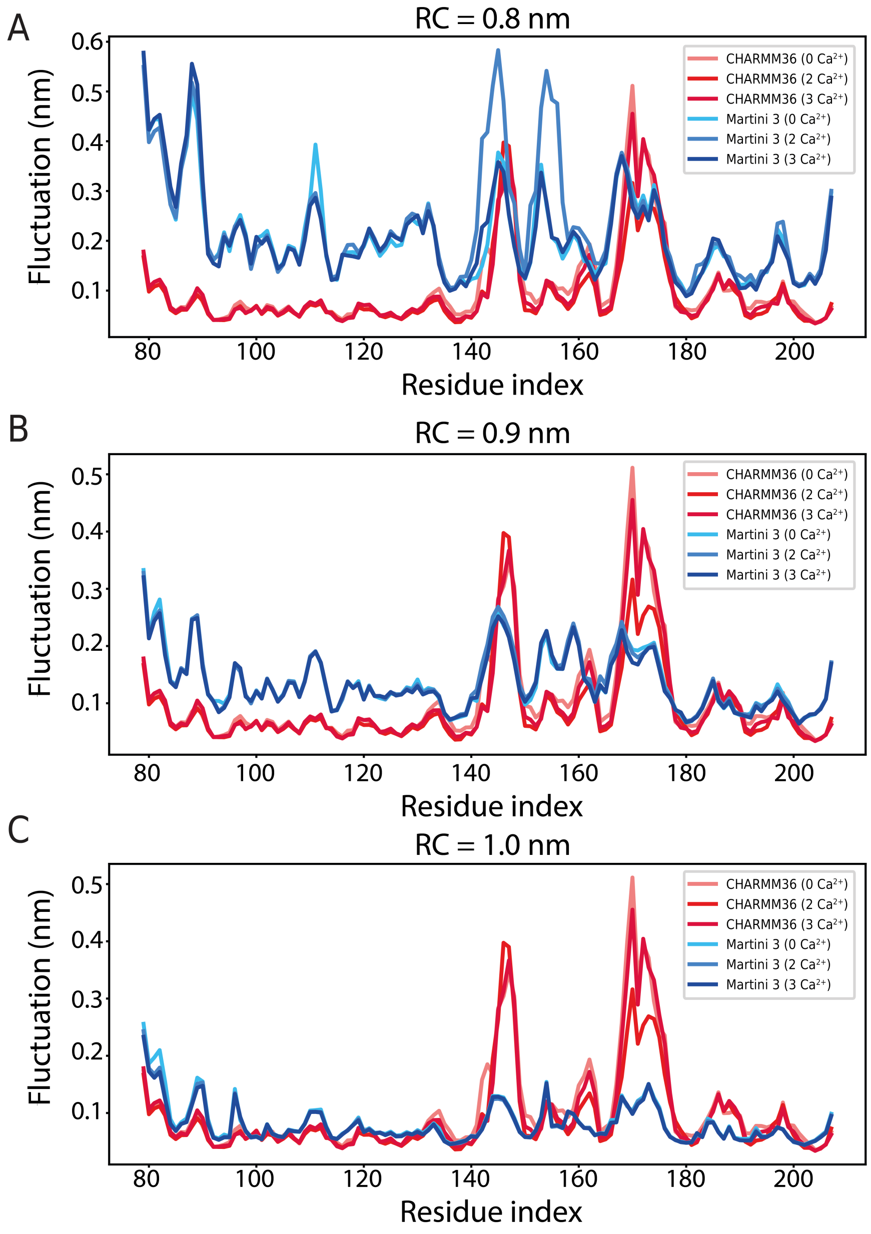


Supporting Figure 3: **Refinement of the elastic network of human Mincle CRD** Comparison of AA (red) and CG (blue) RMSF according to the RC used for the elastic network and the $Ca^{2+}$ concentration.

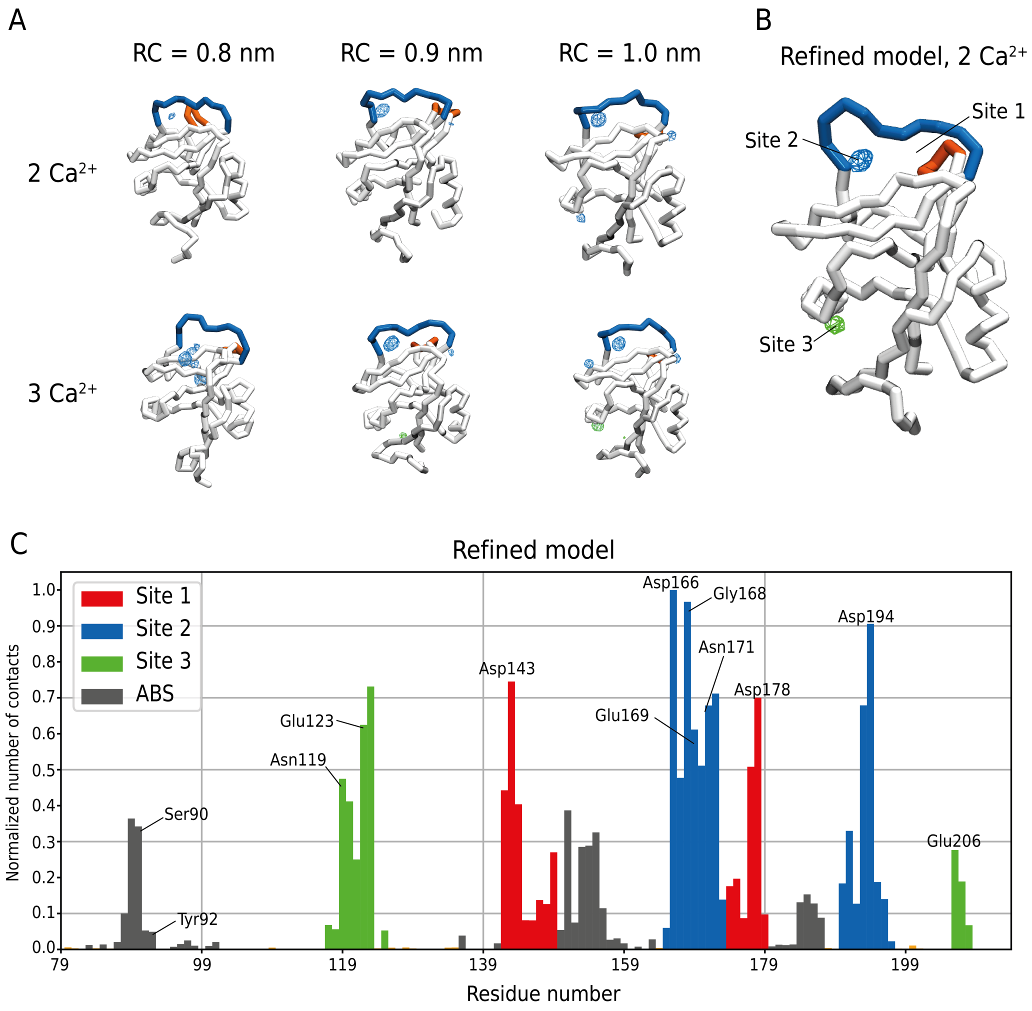


Supporting Figure 4: **$Ca^{2+}$ ions recognition by CG human Mincle CRD** Average position for $Ca^{2+}$ ions for each model with different RC values (A) and for the final refined model (B) (volmap = 0.1). Sites 2 and 3 are shown in blue and green, respectively. The long loop is shown in blue and the short loop in orange. (C) Normalized number of contacts between residues of the refined human Mincle CRD for simulations with 2 and 3 $Ca^{2+}$ ions.

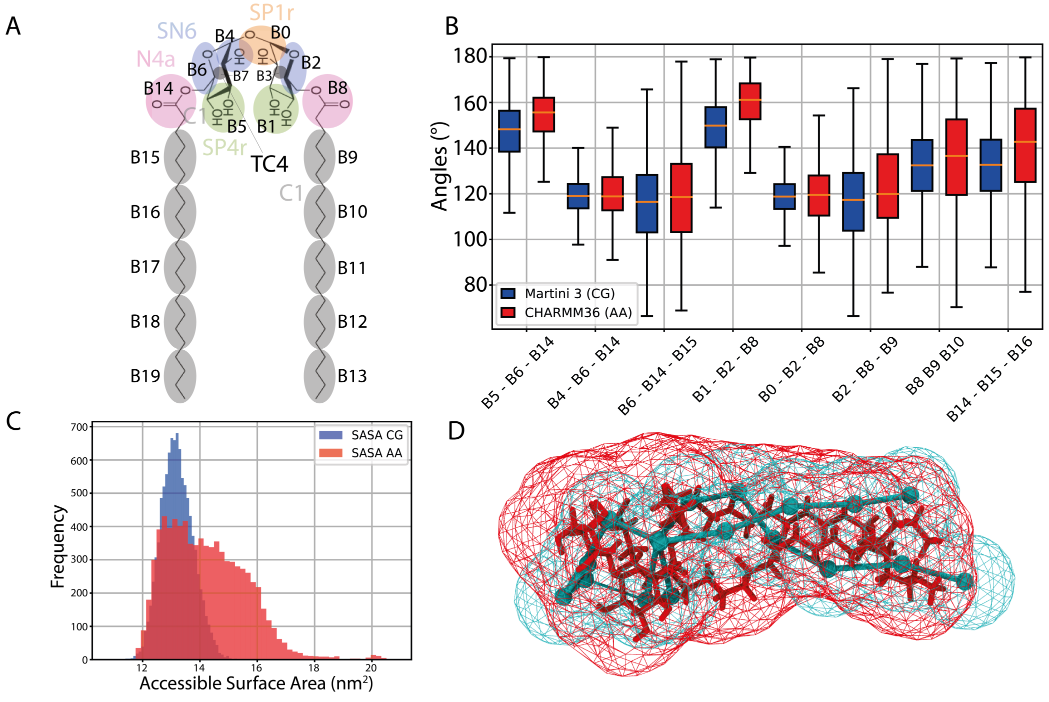


Supporting Figure 5: **Coarse-grained parametrization of TDB** (A) Chemical structure and mapping of TDB. Angles (B) and Solvent Accessible Surface Area (SASA) (C) comparison between all atoms (red) and coarse-grained (blue) simulations. (D) Connolly surface for all-atoms (red) and coarse-grained (blue) TDB.

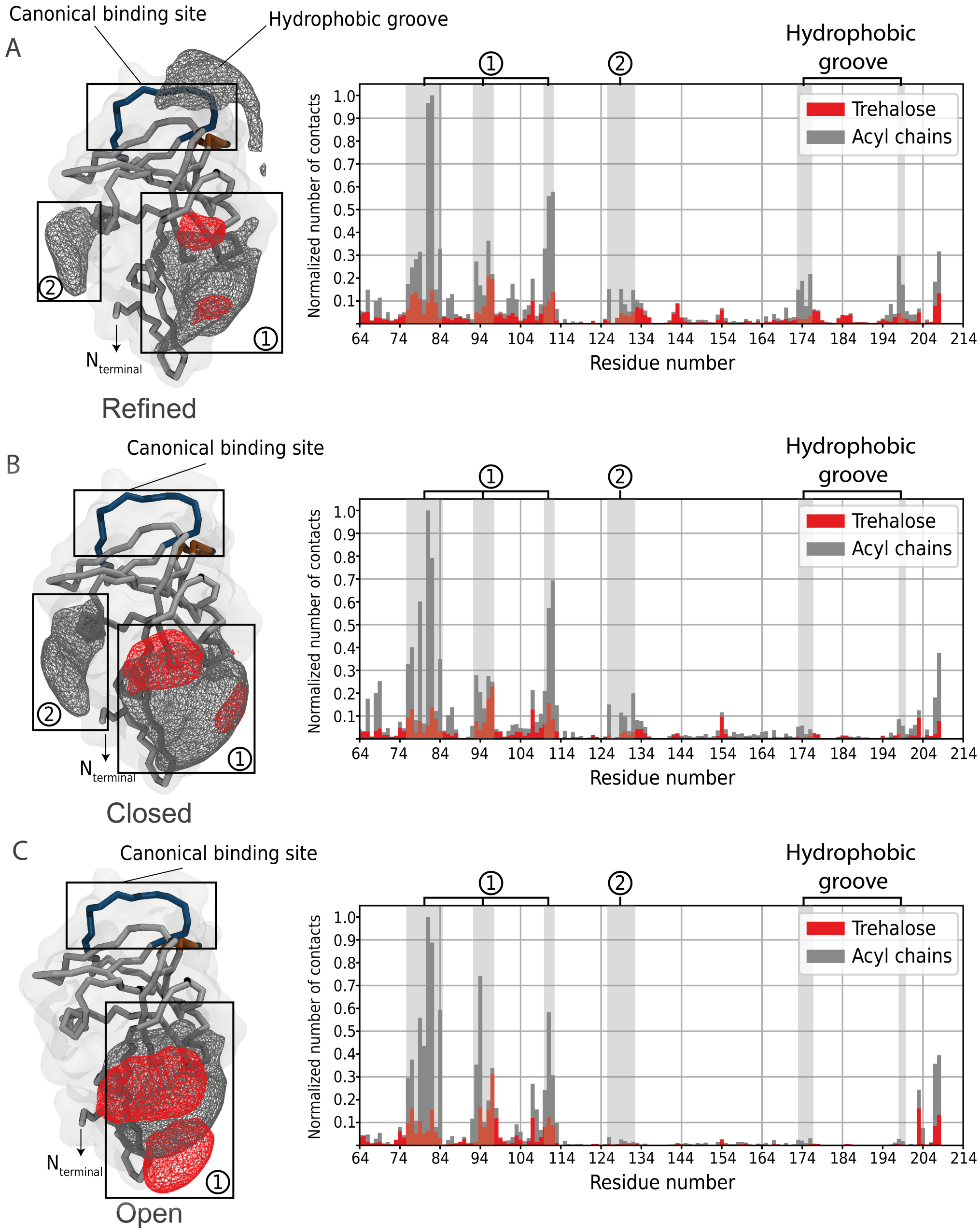


Supporting Figure 6: **Binding of TDB to human Mincle CRD.** Average position of the acyl chains (gray) and trehalose (red) of TDB along 200 μs of unbiased simulation using the refined elastic network (A), the closed conformation (B) or the open conformation (C).

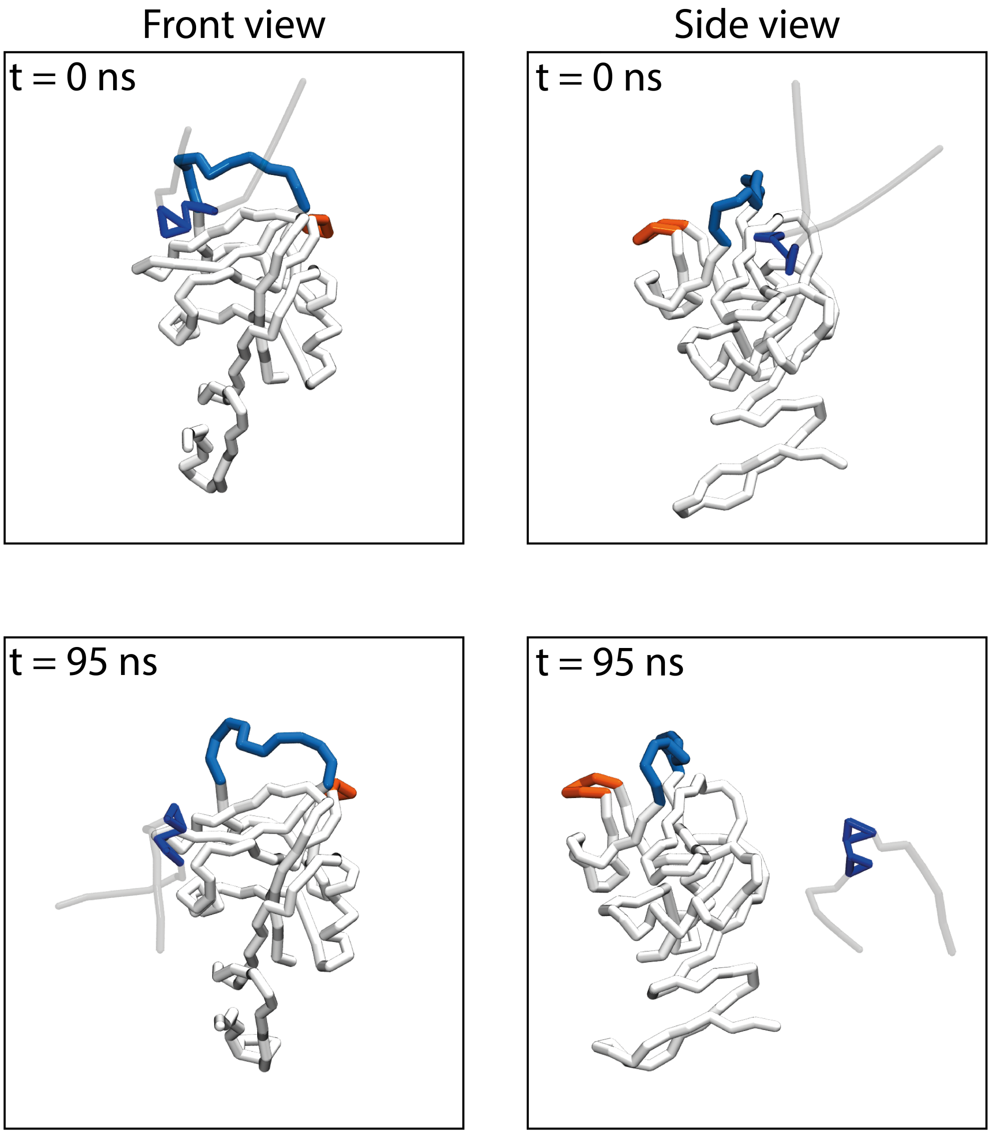


Supporting Figure 7: **Unbinding of trehalose during FEP calculation.** Trehalose with both acyl chains fully decoupled during FEP calculation unbinds after 95 ns of CG simulation. Long and short loops are shown in blue and orange, respectively. The trehalose moiety is shown in blue and decoupled acyl chains in transparent.

Movie 1: **TDB recognition by human Mincle CRD.** Recognition is mediated mainly by residues forming the hydrophobic groove (side chains are shown in white). Long and short loops are shown in blue and orange, respectively. $Ca^{2+}$ ions are represented by red spheres. The trehalose moiety of the TDB is shown in blue and acyl chains in gray.
